# Multimodal mechano-SICM and FRET with stretch for probing cardiomyocyte function

**DOI:** 10.64898/2026.02.11.705278

**Authors:** Benedict Reilly-O’Donnell, Andrew Shevchuk, Julia Gorelik

**Affiliations:** National Heart and Lung Institute, Imperial College London, London, UK; Department of Metabolism, Digestion and Reproduction, Imperial College London, London, UK

**Keywords:** Scanning Ion Conductance Microscopy, Förster Resonance Energy Transfer Microscopy, Cardiomyocyte, Beta-adrenoceptor, Mechanobiology

## Abstract

Cardiac function is dependent upon the ability of cardiomyocytes to adapt their contractions to meet the demands of the body. Increased preload lengthens the sarcomere, altering the efficiency of contraction. The surface topography of cardiomyocytes is distinct from other cell types. T-tubules are key membrane structures which protrude into the cell body, aligned with the edges of the sarcomere. These are key signalling domains which ensure efficient and adaptable excitation-contraction coupling. It has been shown that T-tubules are dynamic structures which deform during the contraction cycle however, how the T-tubule structure adapts to increased preload has not been realised. Here we demonstrate a methodology for the measurement of the surface topography and sub-cellular signalling of isolated adult cardiomyocytes under diastolic stretch. We track individual T-tubule openings, showing that increased load causes them to shift, increase diameter and become stiffer. Future applications of this system include experimental modelling of preload-reducing therapies, for the treatment of acute and chronic heart failure.

## Main Text

### Introduction

The heart adjusts its contraction in a beat-to-beat manner to ensure a good supply of oxygen and nutrients to other tissues. This phenomenon is described by the Frank-Starling relationship^1^, which states that stroke volume will increase in response to increased venous return. Sustained increases in cardiac preload or afterload can disrupt this mechanism of homeostasis, driving a maladaptive response^2^. Preload upon cardiomyocytes can be identified by the sarcomeric length (SL) at the end of diastole. Under normal loading conditions, the diastolic SL remains between 1.90 and 2.20 μm, increased preload lengthens the sarcomere. When SL ≥2.4 μm, the efficiency of contraction reduces^3^.

Increased sarcomeric length of cardiomyocytes, within the physiological range, has been shown to promote Ca^2+^-release through mechanical activation of ion channels^4^ and X-ROS signalling^5^, representing a mechanism of beat-to-beat control of cardiac output. The adrenergic system is a key modulator of cardiac contraction. Recently we reported that β-adrenoceptor-mediated cAMP signalling is regulated by stretch^6^. This study asserted that caveolae structures are key microdomains situated on the cell membrane, which act as signalling centres and membrane reserves-protecting the cell from rupture when stretched^7^. It has also been indicated that T-tubular structures are dynamic throughout the contraction cycle^8^, suggesting that they too could contribute to stretch-regulated β-adrenoceptor signalling. In cases of the failing heart, there is significant remodelling of the T-tubule network^9^, which contributes to altered electrophysiology and aberrant cAMP signalling^10^. It is not clear how alterations in cardiomyocyte preload affect T-tubule structure in the short to medium term and how this may represent a mechanism to modulate β-adrenoceptor signalling.

A limitation of isolated cardiomyocyte studies, is that these cells are not typically subjected to physiological and or pathophysiological preload. The critical link between the form and function of cardiomyocytes is therefore not considered in the majority of studies. This may have caused misunderstanding of associated signalling nanodomains in health and disease, preventing identification of effective treatments. Previous studies which have applied stretch to isolated cardiomyocytes have identified an increase in: sarcoplasmic reticulum Ca^2+^-release^5,11^, ROS production and Ca^2+^-sparks^12^, and recruitment of sarcomeres mediated by titin^13^. How the surface topography of the cardiomyocyte changes under different loads, and how this may modulate cell function, has not yet been identified. Scanning Ion Conductance Microscopy (SICM) generates a topographical map of live cells using a non-contact method, meaning that the technique is particularly advantageous for characterisation of the surface of isolated adult cardiomyocytes^14^.

This study presents an imaging system with the ability to investigate the surface topography and sub-cellular signalling of cardiomyocytes under different loads. We aim to characterise how load affects T-tubule opening structure and β-adrenoceptor signalling.

### Results

The imaging system constructed combines a SICM with micro-controlled stretch apparatus and Förster resonance energy transfer (FRET) fluorescent imaging system (Figure 1A). The pipette-scanning modality of this SICM is particularly advantageous as no sample movement is required (X, Y and Z movement is performed by the nanopipette). For the stretch protocol, isolated cardiomyocytes were plated onto a poly-HEMA coated glass-bottom dish as previously established^15^. Borosilicate glass micro-rods (c.10 μm diameter) were coated in Matrigel and manually positioned on the cell surface. To achieve the desired stretch of the cardiomyocyte, we programmed movement of a single pulling rod (Figure 1B). The cell was stretched as a percentage of the distance between the pulling rods (Figure 1C). Stretch of cardiomyocytes was verified by measurement of SL using “SarcOptiM”^16^.

**Figure 1.**
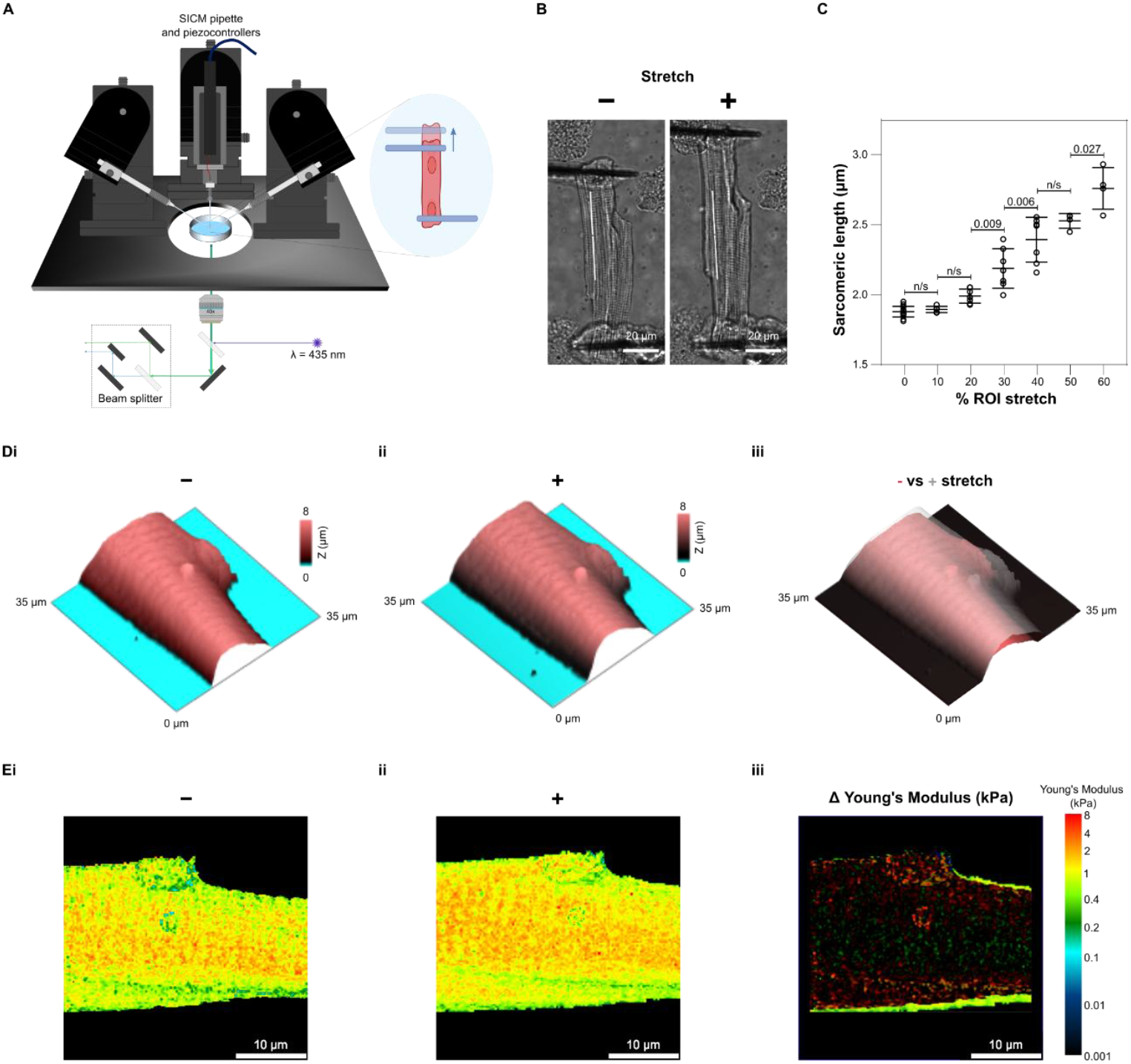
Design of stretch-SICM-FRET system. **A** Illustration of stretch-SICM-FRET system. Two micromanipulators control pulling rods, with a third mounted with the XYZ piezos of the SICM. A FRET light source and beam spitter are mounted underneath. **B** Representative bright field images of adult rat cardiomyocytes pre/post stretch. **C** Sarcomeric length (as determined by SarcOptim^16^) vs % region of interest stretch. n_cells_= 4-14. One-way ANOVA with Tukey’s post-hoc test. Data plotted as mean ±SD **D** Representative 35x35 μm SICM images pre (**Di**) and post (**Dii**) stretch. **Diii** Overlay of images presented in Di (red) & Dii (grey). **E** Representative Young’s modulus maps of SICM images presented in D, pre (**Ei**) and post (**Eii**) stretch. (Eiii) Shift in Young’s modulus, due to stretch, between correlated cell areas.

Repeat scans of the same cardiomyocyte pre (Figure 1Di) and post (Figure 1Dii) stretch identified shifts in membrane features following the direction of stretch. It is possible to track membrane features using this method, allowing for direct comparisons throughout the stretch protocol (Figure 1Diii). The transverse Young’s modulus of the cells was also simultaneously measured by the SICM nanopipette using the intrinsic interaction between the SICM nanopipette and the sample^17^ (Figure 1E).

High resolution 15x15 μm scans (117 nm/px resolution) of the cardiomyocyte clearly displayed surface features, such as T-tubule openings, pre and post stretch (Figure 2A). Surface features could be tracked as load was increased (Figure 2B). It was found that there was no change in the Z-groove ratio (Figure 2Ci) or Z-groove distance (Figure 2Cii) of cardiomyocytes when SL is in the range 1.88-2.75 μm. The diameter of T-tubule openings increased as SL was extended (Figure 2Di). It was not possible to observe any change in the T-tubule depth (Figure 2Dii), however this is likely due to interaction of the nanopipette with the tubule walls, representing a technical limitation of the system. The transverse Young’s modulus of both T-tubule (Figure 2Ei) and crest (Figure 2Eii) regions was increased when cells were stretched beyond SL ≥2.40 μm, similar to previous findings^18^. T-tubules shifted in the same direction as the applied stretch (Figure 2F). These data indicate that, under increased load, T-tubule openings widen and shift without any change to the regularity of the cell surface. Stretching of the cell also increases the transverse Young’s modulus in both the T-tubule and crest regions to a comparable degree.

**Figure 2.**
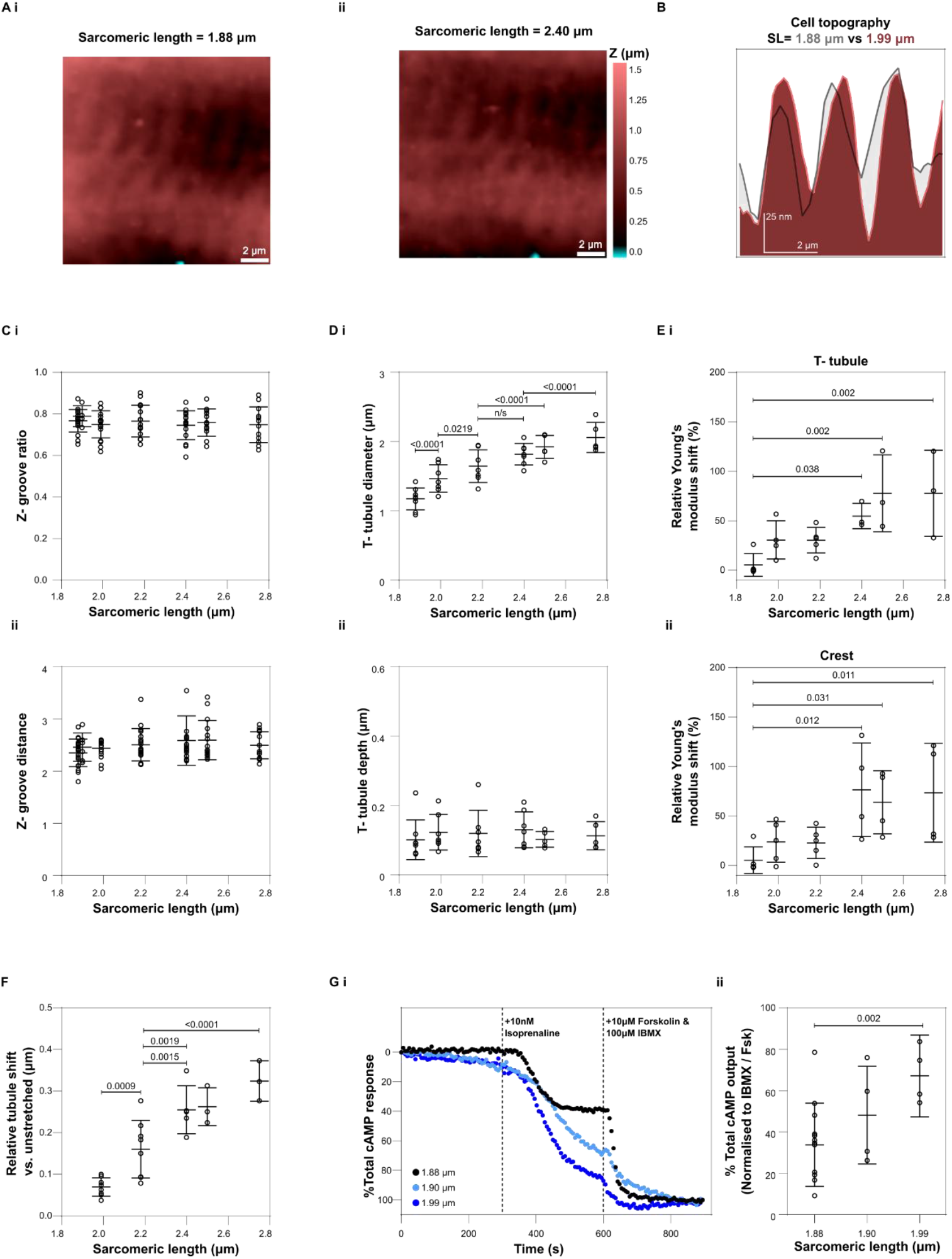
Changes in surface topography and response to β-adrenoceptor stimulation in stretched adult cardiomyocytes. **A** Representative 15x15 μm repeat SICM images of the same cell at two sarcomeric lengths **Ai** SL= 1.88 μm, **Aii** SL= 2.40 μm. **B** Representative line-scan of SICM images of the same cell at two sarcomeric lengths 1.88 μm (red) and 1.99 μm (grey). **Ci** Z-groove ratio of cardiomyocytes at different levels of stretch, calculated from SICM images. n_cells_= 4-17. One-way ANOVA with Tukey’s post-hoc test. **Cii** Z-groove distance of cardiomyocytes at different levels of stretch, calculated from SICM images. n_cells_= 4-17. Kruskal-Wallis followed by Dunn’s multiple comparisons. **D** T-tubule diameter (**Di**) and depth (**Dii**) under longitudinal stretch. n_cells_= 5-8, n_tubules/cell_= 4-14. Nested one-way ANOVA with Tukey’s post-hoc test. **E** Relative Young’s modulus shift in T-tubule (**Ei**) and crest (**Eii**) regions. n_cells_= 3-5, n_ROI/cell_= 3-9. Nested one-way ANOVA with Tukey’s post-hoc test. **F** Relative T-tubule shift (vs. reference T-tubule at SL= 1.88 μm). n_cells_= 3-8, n_tubules/cell_= 4-9. Nested one-way ANOVA with Tukey’s post-hoc test. **G** cAMP response of cells to 10 nM isoprenaline. **Gi** Representative FRET traces of cells expressing EPAC1-cAMPs sensor, normalised to saturator response (100 μM IBMX and 10 μM Forskolin). **Gii** Summary of % total cAMP output in response to 10 nM Isoprenaline. n_animals_= 4-12, n_cells_= 4-17. Nested one-way ANOVA with Tukey’s post-hoc test. All data are plotted as mean ±SD.

To test the responsiveness of the cardiomyocyte to β-adrenoceptor stimulation under different loads, cells were stretched for 5 min and then treated with 10 nM Isoprenaline (Figure 2Gi). Prior to experiments, cells were cultured for 48 hr and infected with the FRET construct EPAC1-cAMPs. The FRET signal produced in response to 10 nM Isoprenaline was normalised to the signal-saturating 100 μM IBMX and 10 μM Forskolin. An increase of cardiomyocyte load to SL= 1.99 μm, caused a significant increase in cAMP output in response to 10 nM Isoprenaline. This correlates with our previous study, which found that hypoosmotic stretch of isolated cardiomyocytes caused an increased cAMP response to β_2_-adrenoceptor stimulation^6^.

### Discussion

The SICM-based system is versatile, with the possibility to also perform nanobiopsies^19^, single-channel patch clamp^20^, and correlative imaging with super-resolution light microscopy^21^. Cell functionality can be further explored using alternate FRET sensors and localised application of drugs^22^, allowing currently unseen characterisation of cardiomyocyte function under load.

The increased diameter of T-tubule openings under load is particularly intriguing, indicating that tubular content exchange may increase in cases of high cardiomyocyte stretch. This may be considered a cardioprotective mechanism by preventing localised depletion of Ca^2+^ and K^+^, promoting stronger contraction through increased entry of Ca^2+^ via the L-type calcium channel and shorter action potential duration.

These data provide the first evidence that the multimodal stretch-mechano-SICM-FRET system has the capability to interrogate the effect of load upon both the cell surface and sub-cellular signalling of adult cardiomyocytes. It is known that that the surface topography of isolated human cardiomyocytes is dynamic and altered in disease^9^. Reducing load in such cells can cause the recovery of some cell features e.g. cell size, but does not fully recover the underlying function^23^. This system could be particularly beneficial for the study of heart failure and its treatment. In particular, preload-reducing therapies, which are commonly prescribed in cases of acute and chronic heart failure^24^.

### Materials and Methods

All animal experiments were carried out in accordance with the United Kingdom Home Office Animals (Scientific Procedures) Act 1986, Amendment Regulations 2012, incorporating the EU Directive 2010/63/EU.

Cardiomyocytes were isolated from the left ventricle of adult male Sprague Dawley rats using the Langendorff perfusion method. For FRET experiments, cells were cultured for 48 hr and infected with the FRET construct EPAC1-cAMPs.

For the stretch protocol, isolated cardiomyocytes were plated on a poly-HEMA coated glass-bottom dish. Borosilicate glass micro-rods (c.10 μm diameter) were coated in Matrigel and manually positioned on the cell surface using a micromanipulator. Cells were stretched using programmed movement of the micromanipulator.

#### Isolation of adult rat cardiomyocytes

All animal experiments were carried out in accordance with the United Kingdom Home Office Animals (Scientific Procedures) Act 1986, Amendment Regulations 2012, incorporating the EU Directive 2010/63/EU.

Cardiomyocytes were isolated from the left ventricle of adult male Sprague Dawley rats (200-500g) using the Langendorff perfusion method as previously described^25^. Animals were anaesthetised with 5% isoflurane and euthanised by cervical dislocation.

#### Manufacture of pulling rods

Pulling rods were formed from borosilicate glass capillaries (BF100-50-7.5, WPI) which were pulled into a 1.5 mm long,10 μm diameter, tip using a Sutter P-2000 puller (Heat-325, Filament-4, Velocity-30, Delay-225 and Pull-130). The rods were bent to a 135° angle using a microforge heating element (Narishige). Bent rods were then rolled in borosilicate glass powder, which formed a thin layer on the rod surface. The rods were then incubated with 4.8 μg/ml Matrigel (Corning) on a rocker (60 RPM, 30 min, RT). Matrigel coating was confirmed by visual inspection. Pulling rods were then stored in PBS, pH= 7.2 at 4 °C until use.

#### Cardiomyocyte attachment and stretch

Isolated cardiomyocytes (c. 5 000 cells) were plated on a 35 mm, poly-HEMA coated, glass-bottom dish^15^ in a high Potassium buffer (120 mM K-gluconate, 25 mM KCl, 2 mM MgCl_2_, 1 mM CaCl_2_, 2 mM EGTA, 10 mM Glucose, 10 mM HEPES. pH= 7.4). Selected cardiomyocytes were then aligned with micromanipulators, to ensure perpendicular attachment of pulling rods to cell. Pulling rods were lowered over each end of the cardiomyocyte until a small deflection was observed in the membrane. The pulling rods were left in position for approximately 5 minutes before lifting the cell from the dish bottom. Static longitudinal stretch was applied by programmed movement of a pulling rod, using MP-285 micromanipulator and software (Sutter). Sarcomere length was monitored using the ImageJ plugin “SarcOptiM”^16^. Cells showed no signs of damage to the membrane in response to stretch.

#### Hopping-mode SICM

Topographical surface images were captured using a scanning ion conductance microscope (SICM) as previously described^26^ the system was controlled using ICAPPIC Universal Controller and Piezo Control System with HPICM Scanner software (ICAPPIC Ltd.). Briefly, an image is obtained using a high-resistance pipette (c. 150 MOhm) attached to X/Y and Z piezos. The current flow through the pipette tip is reduced when it approaches a surface (in the Z-plane). The XYZ position of the pipette, when the current dropped to a pre-defined ‘set point’, was logged before it being withdrawn and ‘hopped’ to the next position. Each XYZ position corresponded to a single pixel of the completed image.

#### SICM image analysis

Z-groove ratio and Z-groove distance were determined using previously described methods^14,22^. T-tubule opening parameters (diameter, depth and shift) were analysed using Gwyddion 2.63^27^. Transverse Young’s modulus was determined using quantitative nanomechanical mapping (QNM) as previously established^17^. QMN was analysed using SICM Image Viewer software (ICAPPIC Ltd.)

#### SICM Detection of intracellular cAMP responses by FRET microscopy

Isolated adult rat cardiomyocytes were plated on (0.3 mg/ml) laminin-coated 35mm dishes. Cells were cultured for 48 hrs in MEM medium and infected with the FRET construct EPAC1-cAMPs by adenoviral delivery^28^. Prior to use, cardiomyocytes were detached from the culture dish using gentle agitation. The cell suspension was centrifuged at 150 xg to pellet the cardiomyocytes. Cells were resuspended in high potassium buffer and plated into a poly-HEMA coated glass-bottom dish. Cells were attached to pulling rods and stretched as detailed above. Cells were illuminated at 435 nm using an LED light source (pE-4000, CooLED). Emitted light was split into donor (CFP) and acceptor (YFP) channels using a beam splitter (Optosplit II, Cairn) fitted with a dichroic mirror (59022bs, Chroma). Images were acquired every 6 seconds with an exposure time of 50 ms.

Non-selective β-adrenoceptor stimulation was achieved by application of 10 nM Isoprenaline. A combination of 100 μM IBMX and 10 μM Forskolin was added to achieve maximal cAMP production within the cell, acting as a signal saturator.

#### FRET microscopy data analysis

The pixel intensity of CFP and YFP images were analysed, post-acquisition, with ImageJ. The FRET ratio (YFP/CFP) was normalised to baseline and calculated. Total cAMP output was calculated as previously described^29^.

#### Statistical analysis

All data sets were tested for normality using the Shapiro-Wilk test. To avoid pseudoreplication we used hierarchical techniques where appropriate, as previously discussed^30^. Significance testing performed is noted in the figure legends. All statistical analysis was performed using GraphPad Prism 8. Data are presented as individual values and mean ±SD.

## Acknowledgments

We thank Jun Woo (Andy) Jang for their assistance in FRET recordings, technical support of Petr Gorelkin, Alexey Yakovlev, and ICAPPIC. We also thank the Imperial College Advanced Hackspace for their assistance in assembling the system and Department of Physics Mechanical Instrumentation Workshop for manufacturing SICM parts.

This study was a collaboration as part of the Cellular Mechanosensing and Functional Microscopy Centre at Imperial College London.

This work was funded by the BHF (RG/F/22/110081). AS acknowledges support of EPSRC (EP/W012219/1).

## Author contributions

BRO performed experiments and wrote the manuscript. AS assisted with SICM imaging. JG supervised the project. All authors edited the manuscript.

